# Novel dynamics of human-carnivore interactions linked to the arrival of *H. sapiens* in Europe

**DOI:** 10.1101/2025.07.31.667895

**Authors:** Vidal-Cordasco Marco, Marín-Arroyo Ana B.

## Abstract

Upon the arrival of *H. sapiens* in Europe, the abundance and diversity of secondary consumers progressively diminished. The factors contributing to this increased human pressure and its potential association with Neanderthal extinction remain unknown. This study identifies biotic and abiotic effects on the structure and assembly of secondary consumers at the European scale during Marine Isotope Stage 3 by integrating analyses of their geographic ranges, co-occurrence patterns, and generalized mixed models. Results show that during the replacement of Neanderthals by *Homo sapiens*, the range of secondary consumers contracted and their co-occurrence frequency increased, leading to new intra-guild interaction dynamics. Additionally, *H. sapiens* occupied a larger portion of the secondary consumers’ fundamental niche. Climate change, the demographic decline of keystone species, and the broader niche breadth of *H. sapiens* reduced the interconnectivity of the co-occurrence network among secondary consumers, shaping novel dynamics of human-carnivore interactions in Europe.

---

Human-carnivore interactions have always held pivotal importance in the study of human evolution. With the incorporation of meat into the *Homo* genus diet, hominins became members of the secondary consumer guild^1^. Myriad debates have emerged regarding the quantity, frequency, and importance of animal protein in their diets^2,3^, the primary or secondary access to the carcasses^4^, and the extent of the subsequent inter-specific competition^5–7^. Due to the dynamic nature of the *Homo* subsistence strategies, the human-carnivore interactions varied across different temporal and spatial contexts^8–10^. A substantial change in this long path of human-carnivore interactions appeared with *H. sapiens*^11,12^.

During the Middle Pleistocene, *H. neanderthalensis* evolved within a rich secondary consumer guild that remained relatively stable in Europe^13^. Archaeozoological evidence shows that Neanderthals competed to some extent for space and resources with bears^14,15^, cave lions^16^ and hyaenas^17–19^. However, when *H. sapiens* replaced Neanderthals, the number of carnivore bones bearing cut marks related to disarticulation and defleshing activities, or used as ornaments, increased meaningfully^20–23^. The high frequency of carnivore bones bearing anthropic modifications, along with the subsequent reduction in carnivore abundance and diversity throughout the Upper Palaeolithic, is frequently interpreted as evidence of increased human pressure on secondary consumers ^24–28^. However, recent studies suggest that these intensified human-carnivore interactions should not be regarded exclusively as instances of competitive exclusion ^29^. Instead, they also involved elements of mutualism, facilitation, or commensalism^30–32^.

There are two main forms of competition: exploitation competition, where species indirectly diminish resources for other species; and interference competition, where one species directly impedes another species’ access to a resource or territory. Whereas interference competition, and consequently avoidance behaviors, are frequently observed between carnivores with similar body sizes and niche overlaps^33,34^, exploitation competition affects all secondary consumer species^35^. However, the intensity of exploitation competition can be mitigated by niche partitioning. Low niche (i.e., spectrum of ecological conditions) and spatial (i.e., geographic range) overlaps between secondary consumers create competition refuges and promote coexistence^33,36^. In contrast, high niche and spatial overlaps increase encounter rates and co-occurrence frequency, leading to higher intra-guild competition^37,38^. Accordingly, the spatial distribution of species arriving in a new ecosystem is commonly assessed to evaluate ecological risk, as it can impact spatial niche partitioning among species. Did the arrival of *H. sapiens* in Europe disrupt the spatial niche partitioning of the secondary consumer guild? To answer this question, the first hypothesis being tested in this study (H_1_) is whether *H. sapiens* occupied a larger portion of the secondary consumers’ fundamental niche during the MIS3 in comparison to Neanderthals.

Biotic interactions between secondary consumers are more complex than bidirectional competitive pressures. Interspecific interactions can be classified as positive, negative or neutral. Positive associations (i.e. aggregations) between sympatric species may arise from mutualistic or facilitative interactions (e.g. kleptoparasitism). Conversely, negative association (i.e. segregations) is often the result of avoidance strategies to alleviate both interference and exploitation competition. Nevertheless, both aggregations and segregations may also emerge due to differences in species’ environmental preferences, spatial distribution and range overlap^39^. Accordingly, the co-occurrence patterns between secondary consumers are contingent upon the species’ abiotic preferences, which define their fundamental niche (i.e., the spectrum of environmental conditions under which a species can survive, defining its potential distribution in the absence of limiting factors such as competition, predation, or diseases), and the biotic interactions that affect their realized niche (i.e., actual suite of conditions under which a species exists).

During the Marine Isotope Stage (MIS) 3 (ca. 60-27 kr BP), abrupt climate changes affected the distribution of several large-mammal species^40^. These climatic fluctuations not only impacted habitat suitability^41^, fragmentation^42^, and isolation^40^, but also altered the abundance and distribution of plant and animal resources^43^, rendering populations more susceptible to bottlenecks and extinctions. In this context of acute climate changes and demographic declines^44^, it is plausible that the spatial range of secondary consumers was constrained, affecting the co-occurrence patterns and the intra-guild interaction dynamics. The second hypothesis this study tests (H_2_) is whether the effects of the MIS3 climate fluctuations on species distributions and spatial overlaps affected the co-occurrence patterns among secondary consumers.

To test both hypotheses (H_1_ and H_2_), we compiled a dataset of 1,970 chronometric dates and 1,561 occurrences of secondary consumer species, comprising carnivores and omnivores, from 373 archaeological and 34 paleontological levels. We ran Bayesian age models to estimate the chronology of each archaeo-paleontological level. Validated and bias-corrected paleoclimate data obtained from the HadCM3 model^45^ were used to build Species Distribution Models (SDM) for: *Ursus spelaeus, Ursus arctos, Panthera spelaea, Panthera pardus, Felis sylvestris, Lynx lynx*, *Lynx pardinus*, *Crocuta crocuta, Canis lupus, Cuon alpinus*, *Vulpes vulpes, Vulpes lagopus, Gulo gulo, Meles meles, Martes martes*, *Mustela erminea*, *Mustela putorius*, *Mustela nivalis, Homo neanderthalensis* and *Homo sapiens* (for details, see “Species Distribution Models” in Methods). A weighted ensemble procedure was used to project the distribution of each species in Europe between 50 and 30 kyr BP. The outputs of the SDMs were converted into favorability values, which represent how well a specific patch aligns with the environmental conditions that are optimal for the species. Habitat favorability ranges between 0 and 1, with values higher than 0.5 corresponding to areas where the probability of presence is higher than expected by chance (for details, see Methods). These projections do not reflect the actual distribution of species, but the evolution of their potential distribution based on their fundamental niche, as described above, and the climatic fluctuation of the MIS3. Habitat favorability was used to evaluate whether, and to what extent, the effects of climate change on niche overlap or differentiation during the MIS3 influenced the observed patterns of species co-occurrence. To this aim, probabilistic co-occurrence analyses assessed the evolution of aggregations and segregations between secondary consumers during the MIS3. Aggregations refer to species that are found together more often than would be expected by chance, while segregations refer to species that are found together less frequently than expected (for details, see “Co-occurrence analyses” in Methods). Lastly, we carried out generalized mixed models to determine whether niche breadth and spatial overlap between secondary consumers affected the observed co-occurrence patterns throughout the MIS3.

## Results

### Spatial distribution range

All SDMs included in this study reached a good predictive capability, with the exception of Generalized Linear Model for *Mustela nivalis* (Supplementary Table 1). Likewise, the five-fold spatial block cross-validation supports the transferability of the model predictions (Supplementary Fig. 1), with the exception of *Mustela nivalis*, *Mustela putorious* and *Lynx pardinus*. Therefore, these three species were excluded in further SDM analyses. The remaining species, for which at least one SDM algorithm achieved an AUC>0.7 in the five-fold spatial block cross-validation (Supplementary Fig. 1), were retained for subsequent analyses. Generalized Linear Models and Bayesian Additive Regression Trees tended to produce the largest extent of favorable areas, while Generalized Additive Models and Maximum Entropy Models projected the smallest (Supplementary Fig. 2). Despite some variation, the evolution of each species’ potential geographic range during MIS 3 remained broadly consistent across most models, with the weighted ensemble projections capturing these common trends (Supplementary Fig. 2). The patterns of variable importance across species show that mean annual temperature, temperature seasonality and mean annual precipitation were the most influential climate variables, whereas rugosity and altitude were the most influential environmental ones (Extended Data Fig. 1). Among secondary consumer species, *Canis lupus, H. sapiens, Lynx lynx, Vulpes vulpes* and *Ursus arctos* had the largest potential geographic ranges. Conversely, *Cuon alpinus*, *Martes martes*, *Panthera pardus* and *Homo neanderthalensis* had, on average, the most restricted geographic ranges (Supplementary Table 2).

Levin’s niche breadth indexes are significantly higher for *H. sapiens* than for *H. neanderthalensis* (Extended Data Fig. 2). Therefore, *H. sapiens* occurred under higher environmental heterogeneity than Neanderthals (i.e., broader niche breadth). During the MIS3, Neanderthals exhibited, on average, a 52.43% smaller potential geographic range compared to *H. sapiens* (Fig. 1). This implies that *Homo sapiens* had the potential to expand over an additional 1,326,052 km² in Europe (Supplementary Table 2), particularly in more septentrional latitudes (Fig. 1).

**Figure 1.**
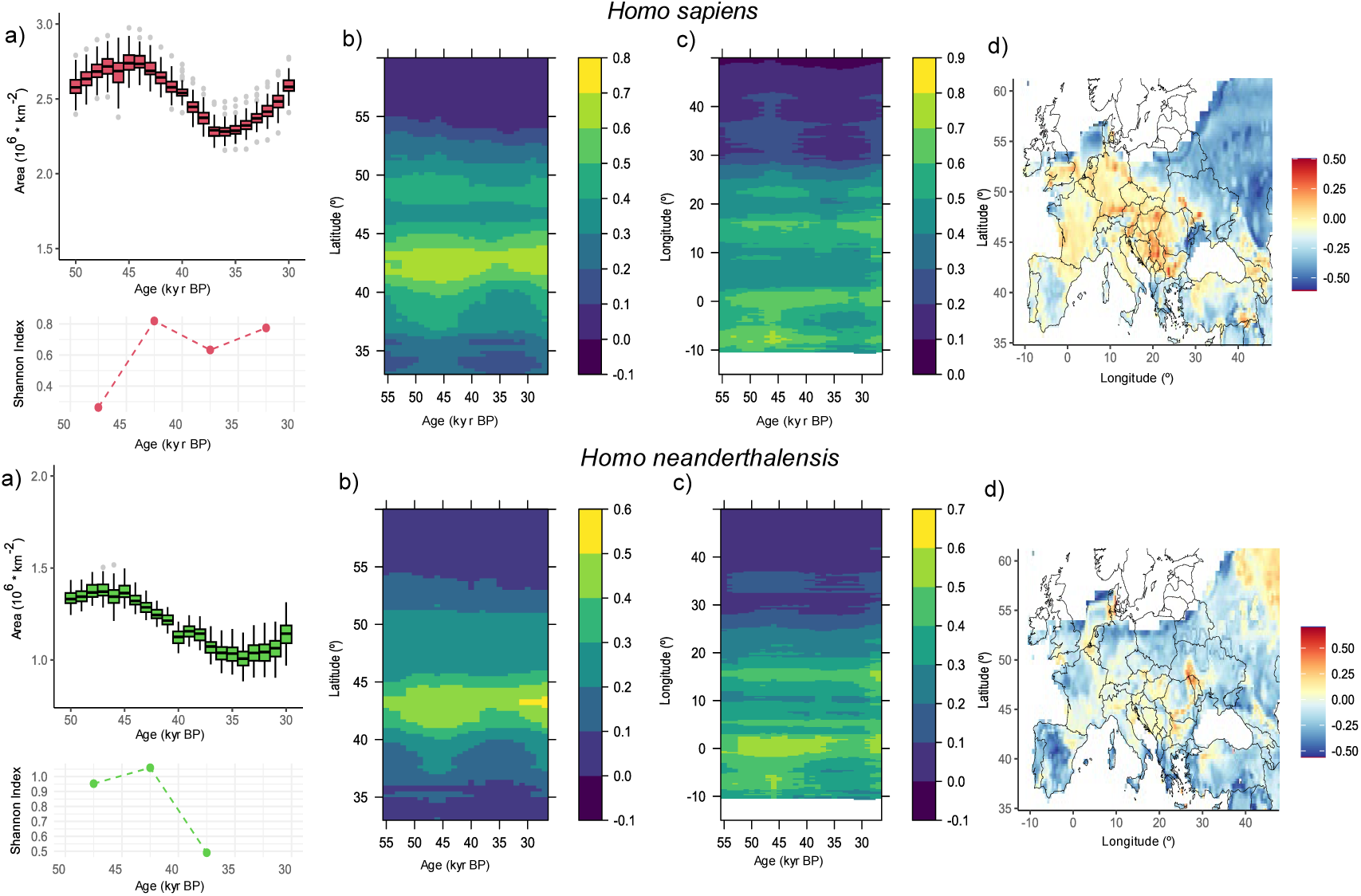
Favorable area for Hominidae species. **a**, Temporal evolution of the geographic extension of favorable areas. Box and whiskers plots represent the interquartile range obtained after repeating the analyses 100 times for each species to account for age uncertainty. Below, the Shannon index reflects the diversity of co-occurring secondary consumer species in bins of 5 k years between 50 and 30 kyr BP. **b,** latitude-time Hovmoeller diagram showing the spatial distribution of the habitat favorability. **c**, longitude-time Hovmoeller diagram showing the spatial distribution of habitat favorability. **d**, percentage difference in habitat favorability between the period with greater extension and stability of favorable areas (50-45 kya) and the period of greatest favorability decline (40-35 kya).

The habitat favorability of *H. neanderthalensis* began to decline by 45 kyr BP, which coincided with an increment in the taxonomic richness of carnivores recovered from the Middle Paleolithic assemblages (i.e., Shannon index) (Fig. 1). Neanderthal habitat favorability reached its lowest level around 35 kyr BP, leading to a loss of 33.8% of their potential geographic range (Fig. 1). The decline in habitat favorability for *H. neanderthalensis* was widespread across Europe and only remained stable in scattered small patches across the continent (Fig. 1d). Between 45 and 35 kyrs BP, the potential range of *H. sapiens* experienced an average contraction of 18.5%, followed by a constant expansion until the end of the MIS3. However, the habitat favorability loss for *H. sapiens* occurred mainly in Eastern Europe, while concurrently it either increased or remained stable in Western Europe. Moreover, in comparison with *H.* n*eanderthalensis*, the potential spatial overlap of *H. sapiens* with all secondary consumer species was, on average, an 82.22% higher (Fig. 2). Therefore, *H. sapiens* might have occupied a larger portion of all secondary consumers’ fundamental niche.

**Figure 2.**
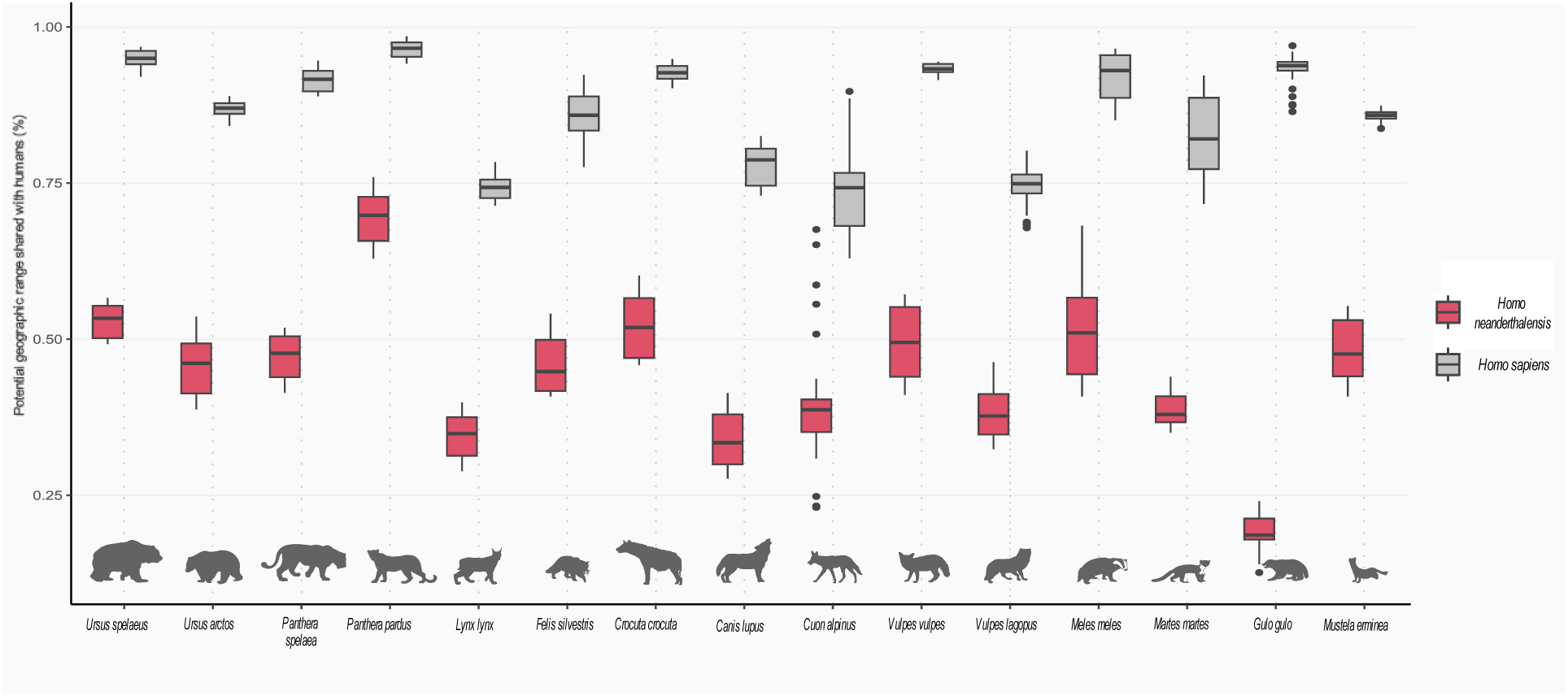
Spatial overlap with humans. Box and whiskers represent the percentage of the spatial range of secondary consumers potentially occupied by humans during the MIS3. Silhouettes sourced from PhyloPic (https://www.phylopic.org/)

Among Felidae, *Lynx lynx* had the largest potential geographic range, followed by *Panthera spelaea, Felis silvestris* and *Panthera pardus* (Extended Data Fig. 3)*. Lynx lynx* was the felid species that exhibited the smallest reduction in habitat favorability during the MIS 3, while *Panthera spelaea* showed the largest decrease. Between 40 and 35 kyr BP, *Panthera spelaea* lost habitat favorability in Eastern and Northern Europe while it remained stable or increased in areas of Western and Southern Europe. In comparison to *Panthera spelaea*, *Panthera pardus* had a substantially narrower latitudinal range and a more fragmented distribution of habitat favorability along the longitudinal range; yet, its habitat favorability remained more stable throughout the MIS3 (Extended Data Fig. 3).

The favorable spatial range of *Ursus arctos* was, on average, 32.1% larger than that for *Ursus spelaeus* (Extended Data Fig. 4). Both species shared a similar spatial pattern of habitat loss between 45 and 35 kyr BP, which probably resulted in population contractions toward the same areas. As the habitat favorability decreased, the Shannon index increased. Accordingly, the onset of the spatial contraction trend coincided with an increase in the diversity of carnivores found along with Ursidae species in the European assemblages. On average, both species shared 1,407,203 km^2^ of favorable areas in Europe, which represents 65.4% of the *U. arctos* and 86.3% of the *U. spelaeus*’ potential geographic range. Therefore, both species had a substantial spatial overlap across Europe, but this overlap represents a larger proportion of the *Ursus spelaeus*’ fundamental niche than it does for *Ursus arctos*.

During the period spanning 50 to 35 kyr BP, *Vulpes vulpes* exhibited a larger potential geographic range compared to *Vulpes lagopus*. However, following this period, the potential geographic range of *Vulpes lagopus* surpassed that of *Vulpes vulpes* in Central and Northern Europe (Extended Data Fig. 5). *Vulpes vulpes* shared, on average, 62% of its potential spatial range with *Vulpes lagopus*, showing the latter a higher habitat favorability in more septentrional latitudes. *Canis lupus* and *Crocuta crocuta* had a widespread distribution, and therefore, a large overlap with most carnivores during the MIS3. *Vulpes vulpes*, *Canis lupus* and *Crocuta crocuta* shared a similar spatial pattern of habitat favorability loss between 45 and 35 kyr BP, followed by a rapid recovery by the end of the MIS3 (Extended Data Fig. 5). As their habitats contracted, the diversity of secondary consumers found alongside these species increased. From 35 kyr BP onwards, the habitat favorability of these three species recovered, and the carnivore diversity in the assemblages declined (Extended Data Fig. 5). Mustelids, like other small carnivores (e.g., *Felis silvestris* or *Vulpes lagopus*), experienced contraction in habitat favorability by 45 kyr BP and again by 40 kyr BP (Extended Data Fig. 6). However, unlike large carnivores, the first contraction was more pronounced than the second, followed by a steady recovery that continued until the end of MIS3 (Extended Data Fig. 6).

Between 45 and 35 kyr BP, all species underwent notable contractions in their habitat favorability and transformations in the co-occurrence frequency with other secondary consumer species. Nevertheless, these transformations were not uniform across all species and did not take place in the same regions. These results show that all secondary consumer species experienced meaningful declines in their habitat favorability during the MIS3 (Extended Data Figs. 3-6). Moreover, the most notable contraction in habitat favorability corresponded with increases in the richness of the secondary consumers in the archaeological assemblages between 45 and 35 kyr BP, coinciding with the replacement of Neanderthals by *H. sapiens* in Europe.

### Co-occurrence patterns

To further explore the transformations in the secondary consumers’ co-occurrence patterns before, during and after the replacement of Neanderthals by *H. sapiens*, we carried out probabilistic co-occurrence analyses spanning from 50 to 30 kya in intervals of 5,000 years (Fig. 3). Thus, we divided the co-occurrence analyses into the following four specific periods based on the observed shifts in both habitat favorability and the replacement of Neanderthals by *H. sapiens*:

1. Phase I (50-45 kyr BP) corresponds to the most stable and favorable habitat conditions for secondary consumers, and Middle Paleolithic techno-complexes associated with Neanderthals were dominant in Europe.
2. Phase II (45-40 kyr BP) marks the onset of a declining trend in habitat favorability for most secondary consumer species, accompanied by the emergence of new techno-complexes attributed to *H. sapiens* and the disappearance of Neanderthal-associated techno-complexes in most European regions.
3. Phase III (40-35 kyr BP) is characterized by the lowest habitat favorability values, coinciding with the extinction of Neanderthals in Europe.
4. During phase IV (35-30 kyr BP), *H. sapiens* becomes the only human species in Europe, and there is a recovery in the habitat favorability for most secondary consumer species.

**Figure 3.**
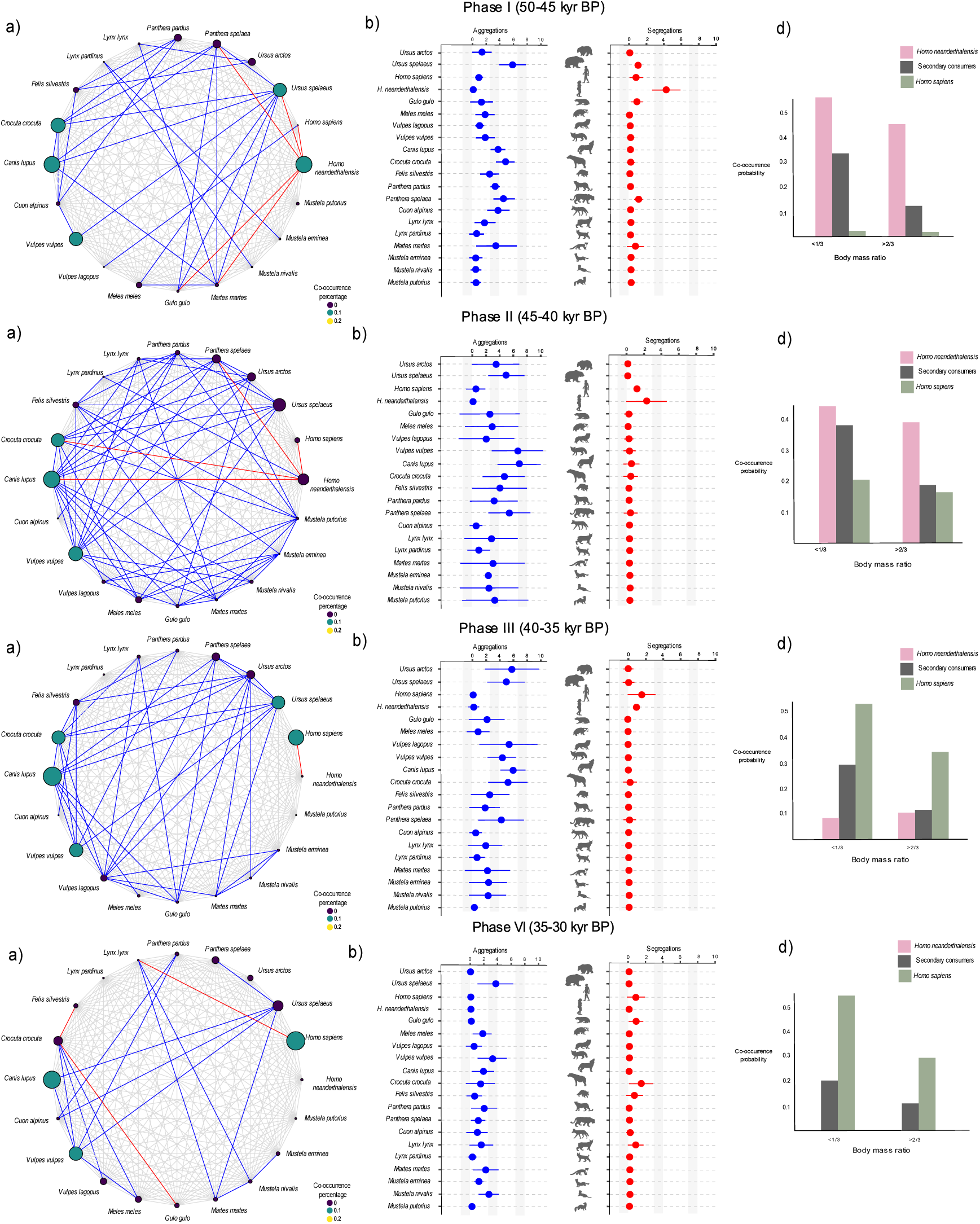
Co-occurrence patterns among secondary consumers. **a**, co-occurrence among secondary consumers between 50 and 30 kya in Europe. Nodes represent species, with their size corresponding to the mean co-occurrence probability. Links represent random (grey), positive (blue) and negative (red) associations between species pairs. **b**, mean number of aggregations obtained from the sensitivity test. Error bars show the 95% CI. **c**, mean number of segregations. Error bars show the 95% CI after the resampling procedure. **d**, Distribution of the co-occurrence probability according to body mass ratio. Silhouettes sourced from PhyloPic (https://www.phylopic.org/)

The species with the highest number of co-occurrences with other secondary consumer species is *Canis lupus*, followed by *Vulpes vulpes* and *Crocuta crocuta* (Table 1). However, throughout the MIS3, there were fluctuations in the number of aggregations and segregations, coupled with transformations in the co-occurrence patterns.

**Table 1.**
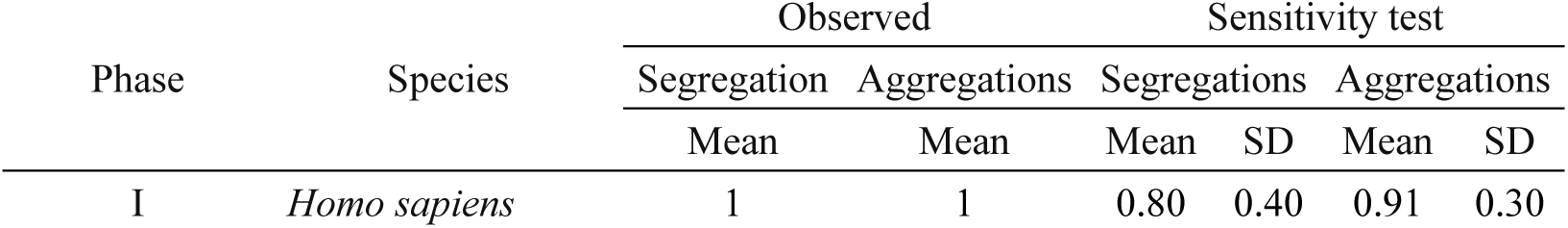
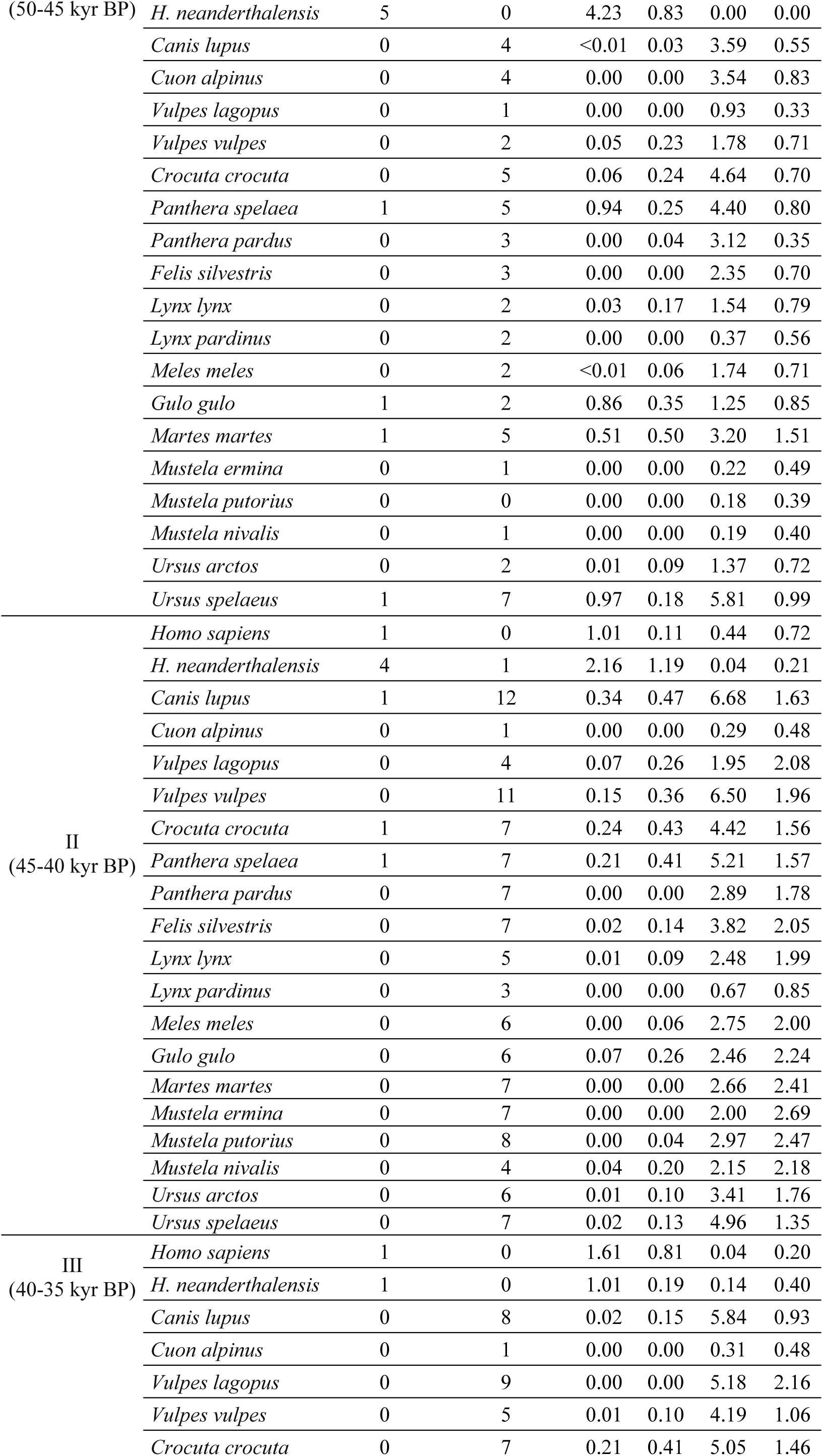
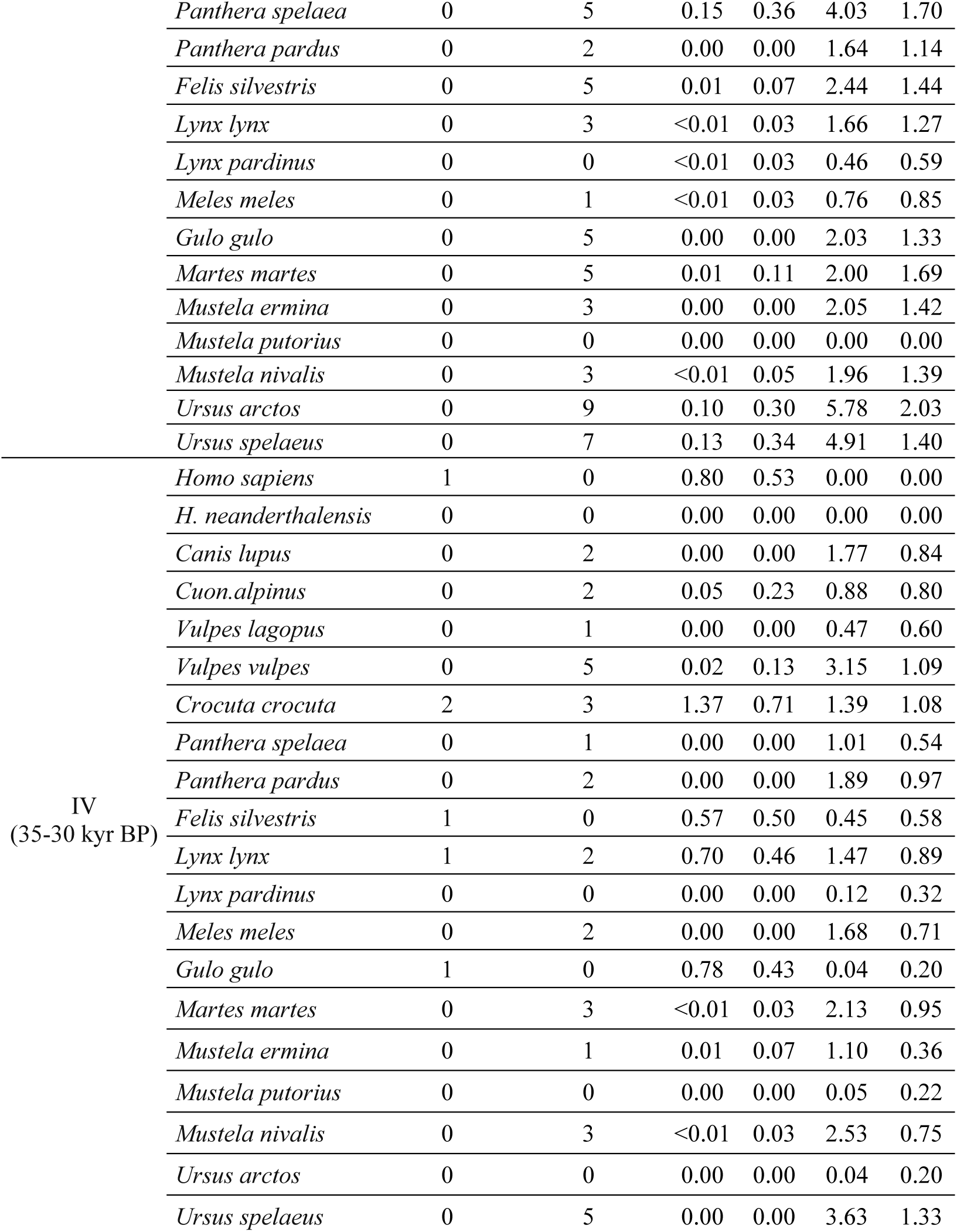
Positive and negative associations. Number of observed segregations and aggregations for each species during the MIS3 phases, compared to the mean number and standard deviation (SD) obtained after conducting 1000 additional co-occurrence analyses from the randomly subsampled assemblages for each phase (for details, see the “co-occurrence analysis” subsection in Methods).

Between 50 and 45 kyr BP, four segregations are observed: between *H. neanderthalnesis* and *Ursus spelaeus,* between *H. neanderthalensis* and *Panthera spelaea*, between Neanderthals and *Martes martes*, and between Neanderthals and *Gulo gulo.* On average, 69.4 % of the favorable areas for *H. neanderthalensis* overlapped with those of U*. spelaeus*, while 78.7% coincided with those of *Panthera spelaea* during this period (Extended Data Fig. 7). In contrast, only 5.1% of the favorable areas for *H. neanderthalensis* overlapped with those of *Gulo gulo*. Therefore, while the segregations between Neanderthals and wolverines can be explained by their limited potential spatial overlap during this period, the segregations with cave bears and cave lions cannot be attributed to different spatial distributions or little spatial overlap.

During the replacement of Neanderthals by *H. sapiens* (45-40 kyr BP), the co-occurrence network in Europe experienced a meaningful transformation, with a two-fold increment in the number of aggregations (Table 1). Over this period, *H. neanderthalensis* is segregated from *Crocuta crocuta*, *Panthera spelaea* and *Canis lupus* (Fig. 3). Moreover, 90.1% of the favorable areas for Neanderthals overlapped with favorable areas for *Crocuta crocuta*, 81.1% with *P. spelaea*, and 99% with *C. lupus* (Extended Data Fig. 7). Therefore, the segregation between Neanderthals and *Crocuta crocuta*, *P. spelaea* and *C. lupus* cannot be be attributed to differences in the distribution of their favorable areas at the European scale between 45 and 40 kyr BP.

Unlike Neanderthals, *H. sapiens* did not exhibited segregations with other secondary consumer species during the MIS3, except for *Lynx lynx* between 35 and 30 kyr BP (Fig. 3). Between 40 and 35 kyr BP, as *H. sapiens* replaced the last Neanderthal populations, the number of aggregations decreased by 32.7%. Yet, with the complete replacement of Neanderthals by *H. sapiens* from 35 to 30 kyr BP, there was a sharper decrease in the number of co-occurrences, with aggregations declining by 59% in Europe (Fig. 3). This pattern, characterized by a substantial increase in the number of aggregations between 45–35 kyr BP followed by a subsequent decline, remains consistent when the co-occurrence analyses are re-run excluding paleontological assemblages (Supplementary Table 3).

Across all periods, the likelihood of co-occurrence was greater for species pairs with a body mass ratio of 1:3 or lower (Fig. 3). Therefore, the co-occurrence probabilities were greater for secondary consumer species pairs with larger body size differences. This effect of body mass ratio on co-occurrence probability applies for Neanderthals-carnivore and *H. sapiens*-carnivore associations (Fig. 3).

### Ecological drivers of species associations

To determine the extent to which the niche breadth (i.e., heterogenity of environmental conditions) of secondary consumers and the overlap of their fundamental niches affected the observed co-occurrence patterns throughout the MIS3, we carried out generalized mixed models. The number of co-occurrences among secondary consumer species exhibit positive correlations with the Levin’s B2 index of niche breadth, the extent of favorable areas, the potential spatial overlap, and the niche overlap indices of Schoener’s D and Warren’s I. Thus, niche breadth, niche overlap, and spatial overlap explain between 15% and 45% of the observed variance in the number of co-occurrences (Table 2).

**Table 2.** Influencing factors on co-occurrence. Outcomes obtained from the generalized mixed models for each predictor (independent) and response (dependent) variable, with the estimated coefficient for each predictor (Estimate), the standard error (Sdt. Error), statistical significance of the coefficient (p-value), marginal correlation coefficient (r^2^) and Akaike Information Criterion (AIC).

| Independent | Dependent |  | Estimate | Std. Error | p-value | r<sup>2</sup> | AIC |
| --- | --- | --- | --- | --- | --- | --- | --- |
| Number of co-occurrences | Levin's B1 | (Intercept) | 6.55 | 2.95 | 0.026 | 0.01 | 2143.8 |
|  |  | Levin's B1 | -2.49 | 3.10 | 0.421 |  |  |
|  | Levin's B2 | (Intercept) | 5.16 | 0.32 | <0.001 | 0.15 | 1881.8 |
|  |  | Levin's B2 | -1.89 | 0.57 | 0.001 |  |  |
|  | Favorable area | (Intercept) | 4.17 | 0.12 | <0.001 | 0.45 | 2078.7 |
|  |  | Favorable area | 0.52 | 0.06 | <0.001 |  |  |
|  | Spatial overlap | (Intercept) | 4.19 | 0.12 | <0.001 | 0.26 | 1876 |
|  |  | Spatial overlap | 0.34 | 0.06 | <0.001 |  |  |
|  | Schoeners'D | (Intercept) | 4.19 | 0.12 | <0.001 | 0.33 | 1894.1 |
|  |  | Schoeners' D | 0.41 | 0.11 | <0.001 |  |  |
|  | Warren's I | (Intercept) | 4.19 | 0.14 | <0.001 | 0.19 | 1899.5 |
|  |  | Warren's I | 0.31 | 0.13 | 0.021 |  |  |
| Aggregations | Levin's B1 | (Intercept) | -0.47 | 2.52 | 0.850 | 0.0.1 | 315.6 |
|  |  | Levin's B1 | 1.98 | 2.65 | 0.456 |  |  |
|  | Levin's B2 | (Intercept) | 1.47 | 0.28 | <0.001 | <0.01 | 311.8 |
|  |  | Levin's B2 | -0.11 | 0.54 | 0.833 |  |  |
|  | Favorable area | (Intercept) | 1.41 | 0.07 | <0.001 | 0.05 | 313 |
|  |  | Favorable area | 0.12 | 0.07 | 0.082 |  |  |
|  | Spatial overlap | (Intercept) | 1.41 | 0.07 | <0.001 | 0.08 | 312.5 |
|  |  | Spatial overlap | 0.15 | 0.07 | 0.037 |  |  |
|  | Schoeners'D | (Intercept) | 1.40 | 0.07 | <0.001 | 0.15 | 310.2 |
|  |  | Schoeners' D | 0.21 | 0.08 | 0.010 |  |  |
|  | Warren's I | (Intercept) | 1.40 | 0.07 | <0.001 | 0.16 | 310.4 |
|  |  | Warren's I | 0.21 | 0.08 | 0.013 |  |  |
| Segregations | Levin's B1 | (Intercept) | 9.20 | 15.85 | 0.562 | 0.02 | 55.5 |
|  |  | Levin's B1 | -9.33 | 16.65 | 0.575 |  |  |
|  | Levin's B2 | (Intercept) | 1.81 | 1.42 | 0.203 | 0.08 | 54.7 |
|  |  | Levin's B2 | -3.046 | 2.917 | 0.296 |  |  |
|  | Favorable area | (Intercept) | 0.32 | 0.23 | 0.161 | 0.06 | 55.1 |
|  |  | Favorable area | -0.19 | 0.22 | 0.372 |  |  |
|  | Spatial overlap | (Intercept) | 0.31 | 0.23 | 0.179 | 0.02 | 55.5 |
|  |  | Spatial overlap | -0.126 | 0.22 | 0.572 |  |  |
|  | Schoeners'D | (Intercept) | 0.31 | 0.24 | 0.196 | 0.02 | 55.6 |
|  |  | Schoeners' D | 0.11 | 0.24 | 0.648 |  |  |
|  | Warren's I | (Intercept) | 0.31 | 0.24 | 0.198 | 0.02 | 55.5 |
|  |  | Warren's I | 0.13 | 0.24 | 0.591 |  |  |

The number of segregations is not significantly correlated with any of the independent variables. Conversely, among the aggregated secondary consumer species, the number of positive associations was significantly higher for those species with higher values of niche overlap (Warren’s I and Schoener’s D) and spatial overlap (Table 2). These results suggest that niche breadth, spatial overlap, and niche overlap significantly affected the observed co-occurrence patterns during the MIS3, highlighting the importance of these factors in shaping ecological interactions among species.

## Discussion

The structure of mammalian communities is shaped by a combination of species’ environmental preferences (i.e., abiotic factors) and interactions among species (i.e., biotic factors)^33,39^. As archaeological and paleontological assemblages encompass large temporal scales, disentangling the effects of biotic and abiotic factors on the observed co-occurrence patterns represents a major challenge. By integrating species distribution models, probabilistic co-occurrence analyses and generalized mixed models, this study highlights the underlying processes of the MIS3 secondary consumers’ community structure and assembly. The obtained results hold implications beyond the comprehension of the intra-guild co-occurrence dynamics among secondary consumers during the Neanderthal demise. They also highlight the effects caused by climate changes and the arrival of our species in Europe, offering insights into broader ecological and evolutionary processes.

Traditionally, human evolutionary studies have explored the distinctive traits of *H. sapiens* to explain why we are the only human species currently on Earth. These studies primarily focused on symbolism^46^, cognition^47^, societal dynamics^48^, and communicative skills^48^. However, with the growing concern over biodiversity loss driven by human activities in recent times, there has been a shift towards examining the ecological impact of *H. sapiens* in our deep past^49,50^. In this context, scholars have stressed the ecological adaptability of our species since the Pleistocene^42,51,52^; yet, the debate ensues regarding the implications of this plasticity. This study provides evidence not only for the wider niche breadth of our species in comparison to Neanderthals, but also for its consequences for the secondary consumers’ guild structure and assembly.

When the presence of *H. sapiens* was scarce and the Middle Paleolithic techno-complexes were dominant on the European continent (50-45 kyr BP), the habitat favorability of secondary consumers remained relatively stable. However, a general habitat favorability loss process started with the severely cold and arid conditions experienced in Europe between 45 and 35 kyr BP^44^, coincident with the rise of modern human settlements in Europe (i.e., techno-complexes associated with *H. sapiens*, for details, see Supplementary Note 1)^53,54^, the emergence of new techno-complexes often referred to as “transitional industries”^55,56^, and the progressive disappearance of techno-complexes assotiated with Neanderthals^43,57,58^. Therefore, the demographic decline of Neanderthals was part of a broader trend of habitat loss for most of the MIS3 secondary consumers^59^ that led to population contractions toward refuge areas of Southern and Western Europe, where carnivore density, richness and intra-guild interaction dynamics experienced substantial changes.

The extent of favorable areas, niche breadth, niche overlap, and spatial overlap among species account for approximately one-third of the variance observed in the number of co-occurrences, and about 15% of the variance observed in the number of aggregations. Accordingly, climate conditions during the Middle to Upper Palaeolithic transition not only contracted the spatial range and affected the overlap extent between secondary consumer species, but also affected their co-occurrence patterns. Thus, habitat favorability contractions between 45 and 35 kya concentrated secondary consumers within smaller and more isolated patches, which led to an increment in the carnivore taxonomic diversity in both archaeological and paleontological assemblages.

The number of observed segregations is not correlated with niche breadth, niche overlap, or spatial overlap among species. Five out of the six observed Neanderthal segregations occurred with secondary consumers that shared more than two-thirds of their spatial distributions. These results suggest that most of the observed segregations between Neanderthals and carnivores cannot be explained by differences in their spatial ranges. Instead, these segregations may be the result of biotic dynamics of avoidance, especially as the potential geographic range of carnivore species contracted in Europe. However, these segregations could also be linked to changes in other aspects of their ecological niches that were not considered in this study, such as shifts in foraging or dietary behaviors that led to niche partitioning, or challenges some species faced in adapting to new ecological conditions, including more fragmented habitats^60^.

Apex predators often exhibit spatial antagonistic associations as they engage in direct competition with other top predators for resources and space^33^. In this connection, the number of segregations in the Middle Paleolithic assemblages was larger than in the Upper Paleolithic assemblages. However, this difference in the number of negative associations does not imply that Neanderthal-carnivore competition was higher than that of *H. sapiens*. On the contrary, the lower number of *H. sapiens* segregations likely reflects their greater spatial overlap with carnivores, and therefore, the ability of our species to occupy a larger portion of the whole secondary consumers’ fundamental niche.

Overlap in habitat use is frequently used to assess the disruptive potential of new species in ecosystems. On this basis, this study shows that *H. sapiens* occupied a larger portion of all secondary consumers’ fundamental niche since its arrival in Europe, potentially limiting spatial niche partitioning among carnivores. The consequences of this higher “invasive”^61^ potential left various evidences in the archaeological record, such as disruptions in carnivore cave-denning behaviors^62,63^, increased number of anthropic marks on carnivore remains^20,22,27^, or declines in carnivore demography and richness.^26,28^ On the other hand, recent studies indicate that, during the Upper Palaeolithic, *H. sapiens* also influenced some carnivores’ feeding behaviors, leading to new synanthropic niches for ravens, foxes or wolves^23,64–66^. Thus, the greater potential of *H. sapiens* to occupy a larger portion of the secondary consumers’ fundamental niche is probably the foundation for these evidences that reflect a crucial role our species in influencing the synecological dynamics of secondary consumers.

Neanderthals co-evolved along and coexisted with a secondary consumer guild that remained relatively stable since the Middle Pleistocene.^13^ Results obtained in this study suggest that this coexistence between Neanderthals and carnivores on a continental scale was upheld stable for a long period by the spatial structuring of the competitive landscape. Thus, on average, 66.3% of the suitable patches for secondary consumers did not overlap with those of Neanderthals. As in present-day ecosystems, intra-guild differences in the spatial distribution of source and sink areas are crucial for the coexistence between secondary consumers. However, the arrival of a new human species and the general habitat contractions due to the MIS3 climate fluctuations disrupted the previous structure and assembly dynamics.

This study contributes to the growing consensus on the combined effects of climate changes and the arrival of *H. sapiens* in Europe during the MIS3 as key factors that generated ecological transformations that led to demographic declines in different mammal species, including Neanderthals^67,68^. The extinction of *H. neanderthalensis* shares some similarities with the demographic decline of other secondary consumers during the MIS3. *H. neanderthalensis* and *Ursus spelaeus* were large-sized secondary consumers that, during the MIS3, shared an important portion of their geographic distribution with phylogenetically closely related species, with which they potentially competed (i.e. *H. sapiens* and *Ursus arctos* respectively). In both cases, the species that went extinct during the MIS3 or shortly thereafter had narrower geographic ranges and more isolated favorable areas due to habitat losses between 45 and 35 kyr BP. This is consistent with the idea that specialists with narrower or more fragmented ranges are at higher risks of extinction.^39,42,69^

Once *H. sapiens* fully replaced Neanderthals (35-30 kyr BP), the habitat favorability of most species started a recovery trend and the co-occurrence network topology became less heterogeneous, characterized by a reduced number of aggregations and segregations. By the end of the MIS3, some keystone species declined demographically (e.g. *Crocuta crocuta*^70^, *Panthera spelaea*^71^, *Canis lupus*^72^) or disappeared in Europe (i.e. *Ursus spelaeus, H. neanderthalensis*) which probably had cascading effects and led to simpler ecological structures and interactions. Therefore, the reduction in the number of aggregations and segregations by the end of the MIS3 probably results from the combined effects of the aforementioned climate-induced shifts in distribution and spatial overlap between secondary consumers, the extinction of keystone species, and the arrival of *H. sapiens*. Based on the data presented here, we hypothesize that species to have likely benefited from these transformations were those capable of occupying the niches of species that experienced demographic contractions (e.g., *U. arctos* benefiting from the decline of *U. spelaeus* or *H. sapiens* from *H. neanderthalensis*). Additionally, smaller carnivores may have directly benefited from *H. sapiens*’ broader niche breadth, potentially gaining access to their leftovers^31,32^, or indirectly through trophic release effects (i.e., increased herbivore biomass availability triggered by the demographic decline of larger carnivores). In any case, determining which species benefited from these transformations is challenging and requires further investigation.

As in present-day terrestrial ecosystems, the number of aggregations throughout the MIS3 was systematically more frequent than segregations, particularly for mesocarnivores^36^. This suggests that biotic interactions between secondary consumers were more complex than simple competitive pressures, encompassing a large number of mutualistic or facilitative relationships. The fact that wolves, foxes, and hyenas are highly flexible feeders, and were the species with the highest number of co-occurrences, underscores the importance of facilitative dynamics among carnivores. This pattern is often attributed to the wide range of feeding behaviors exhibited by these species, which makes spatial aggregations more common than avoidance behaviors^33,73^. In addition, this study shows that the proposed body size ratio of 1:3 observed in present-day ecosystems as an indicator of co-occurrence likelihood^34,35,74^ holds during the MIS3 in Europe. Accordingly, secondary consumer species with greater body mass differences had a higher probability of co-occurring. This is attributed to the tendency in terrestrial communities for species with different body sizes and foraging strategies to engage in interactions that positively impact the success of at least one species ^35,38^.

A number of limitations should be acknowledged in relation to the present findings and their interpretations. First, assemblages associated with the Châtelperronian, Uluzzian, and Szeletian were excluded from the human–carnivore co-occurrence analyses due to ongoing debates about their authorship and their limited representation in the dataset (see Supplementary Note 1 for details). On the other hand, spatial avoidance or aggregation between species can occur at different scales. In this study, we focus on macroecological processes that influenced the spatial distribution and co-occurrence patterns among secondary consumers on a continental scale, so probably fine-scale differences between species in their niches (e.g., differences in temporal activity patterns or diet preferences) contributed to the high number of aggregations observed in this study. Although this study shows a statistically significant effect of niche breadth, spatial range, and niche overlap on the number of aggregations, a substantial amount of variance remains unexplained. A variety of factors, including prey distribution and abundance, can influence the actual distribution and co-occurrence patterns of secondary consumers. Therefore, further research is needed to expand upon these findings and explore additional factors that may have contributed to the co-occurrence transformations during the Middle to Upper Paleolithic transition in Europe.

The sensitivity tests ruled out the effects of sampling biases on the observed co-occurrence patterns and accounted for age and climate uncertainties in the habitat favorability and spatial distributions. Taking into account these factors, this study disentangled some of the factors that influenced the co-occurrence patterns between secondary consumers. Results are robust in showing that climate changes during the Middle to Upper Paleolithic transition contracted the geographic range of most secondary consumers and disrupted the co-occurrence network of secondary consumers in Europe, leading to an increment of aggregations and segregations. This suggests that, between 45 and 35 kyr BP, there was a meaningful increment in the carnivore density and intra-guild interaction dynamics in Western European regions. The subsequent disappearance of keystone secondary consumer species by the end of the MIS3 and the replacement of Neanderthals by a new human species with a 82 %higher spatial overlap with all secondary consumers led to a less interconnected network of co-occurrences. Thus, the novel dynamics of intra-guild interactions among secondary consumers were affected by the arrival of *H. sapiens* and the onset of the Quaternary megafauna extinctions^75,76^. These events likely triggered trophic cascades and disrupted the previous synecological dynamics^77^. The arrival of a new human species, with a larger niche overlap with secondary consumers, further altered these interactions and limited the spatial niche partitioning among carnivores, leading a lower number of aggregations. Thus, this study identifies some of the factors that contributed to shaping novel dynamics of human-carnivore interactions with the arrival of *H. sapiens* in Europe.

## Materials and Methods

### Database

We collected from the literature a dataset of 1,561 occurrences of secondary consumer species from 405 European archaeological and paleontological levels. This database includes 1,970 chronometric dates recovered from MIS3 dated layers, their quality assessment (for details, see the “Chronology” subsection), and information on the stratigraphic context. The database records all the secondary consumer species recovered from each archaeological or paleontological level, and their body mass according to the Phylacine database^78^. In instances where only the genus is identified, this information is recorded; however, only taxonomic identifications at the species level were utilized in the analyses. Therefore, all dated assemblages containing faunal remains, independently of their lithic tools, were used in the carnivores’ SDMs (for details, see Species Distribution Models subsection). As for the humans’ SDMs and the human-carnivore aggregations and segregations analyses, the attribution of archaeological assemblages to *Homo sapiens* or *Homo neanderthalensis* was based on:

a. The integrity of the assemblage. If mixing between cultural layers or unresolved stratigraphic issues were noted in the literature, the corresponding assemblage was excluded as an indicator of the presence of *H. sapiens* or *H. neanderthalensis*. Justifications for these case-specific exclusions are provided in the database.
b. If human remains, genetic, or proteomic evidence identified the specific hominin species associated with an archaeological layer, that species was recorded regardless of whether lithic tools were recovered.
c. When a diagnostic tecno-complex was identified in an archaeological layer and there is strong evidence and consensus among scholars regarding its hominin authorship, we assigned the corresponding species. Accordingly, the Middle Palaeolithic industries, the Mousterian and the Micoquian assemblages were classified as indicative of the presence of *H. neanderthalensis.* The Upper Paleolithic, Neronian^79^, Aurignacian^80^, Gravettian, Uluzzian^81,82^, LRJ^55^, Early and Initial Upper Palaeolithic^83^ techno-complexes were classified as indicative of *H. sapiens.* Therefore, here we distinguish between two levels of attribution: i) the traditional techno-complexes, and ii) the amalgamations of techno-complexes that designate broader periods (e.g., Upper Paleolithic). Justifications for each of these technocomplex-species associations are detailed in Supplementary Note 1. Assemblages associated with the Szeletian (<1% of the assemblages), Uluzzian (1.7% of the assemblages), and Châtelperronian (2.4% of the assemblages) pose challenges due to ongoing debates about their authorships. Neanderthal remains associated with Châtelperronian lithic tools were recovered from Saint-Césaire and Grotte du Renne ^84,85^. Nevetheless, the Châtelperronian-Neanderthal association has been questioned^86^. Additionally, it has been proposed that an ilium recovered from the Châtelperronian layer of the Grotte du Renne may have belonged to *Homo sapiens* ^87^. As for the Uluzzian, two deciduous teeth recovered from Grotta del Cavallo suggest modern human authorship for the Uluzzian ^88,89^, but some authors call the reliability of this association into question ^90^. Lastly, the Szeletian attribution to *H. sapiens* or *H. neanderthalensis* also remains unresolved ^91,92^. Given these uncertainties, as well as the limited representation of the Châtelperronian, Uluzzian and Szeletian assemblages in our study (<5% of total assemblages), we excluded them from the human–carnivore co-occurrence analyses. Yet, as previously noted, these assemblages containing faunal remains were used in the carnivores’ species distribution models (SDMs).

### Chronology

Following previous studies, radiocarbon dates obtained with ultrafiltration, Amino Acid Racemization (AAR), Accelerator Mass Spectrometry (AMS), or Acid-Base-Oxidation-Stepped Combustion (ABOx-SC) protocols were used in the main analyses^43^. Conventional carbon-14 (14C) measurements, as well as chronometric determinations with potential contamination, low collagen yield, or dates obtained from burnt bones, were excluded from the chronological models^43,58^. Radiocarbon dates obtained from charcoal or bone remains were calibrated using the IntCal20 calibration curve^97^, whereas those dates obtained from marine shells were calibrated with the Marine20 calibration curve^98^ with a ΔR of 0^99^. Additionally, OSL/TL and U/Th chronometric determinations were also incorporated into the chronological models by using one sigma error.

Building on previous studies, we used Bayesian age modeling, as it integrates multiple lines of evidence from the archaeological record by incorporating prior knowledge, such as sequence order and instances where the same sample was dated multiple times. We built hierarchical Bayesian age models with the ChronoModel software^100^. In these chronological models, each event represents one date, and multiple chronometric measurements obtained from the same sample were incorporated into the same event^101^. Each phase comprises a group of events recovered from a particular archaeo-paleontological level. Following previous studies, we used the default Markov chain Monte Carlo settings, consisting of three chains, 1000 burn iterations, and 500 batch iterations with a maximum of 20 batches and 100,000 acquisition iterations with a thinning interval of 10^58,101^. The modeled age and its standard deviation were incorporated into the database and used as the chronology for the species pool recovered from each assemblage.

### Climate variables

We used the HadleyCM3 model^45^, obtained from the Pastclim R package^102^. HadleyCM3 is a coupled general climate model with active atmosphere, ocean and sea ice components. The spatial resolution is 0.5° * 0.5°, with the 17 climate and 5 environmental variables at 1000-year intervals.^45^ We selected this paleoclimate model because its bias-corrected values have been recently validated against paleoclimate data derived from pollen-based temperature and precipitations, as well as stable oxygen values from different stalagmites in Europe during the MIS3^43^. This multi-proxy validation showed that the simulated paleoclimate values capture the climate trends during the MIS3 in Europe^43^. The following 21 variables were used: mean annual temperature, temperature seasonality, max. temperature of the warmest month, min. temperature of the coldest month, temperature annual range, mean temperature of wettest quarter, mean temperature of driest quarter, mean temperature of warmest quarter, mean temperature of coldest quarter, mean annual precipitation, precipitation of wettest month, precipitation of driest month, precipitation seasonality, precipitation of wettest quarter, precipitation of driest quarter, precipitation of warmest quarter, precipitation of coldest quarter, net primary productivity, lead area index, altitude over the sea level and rugosity.

### Species Distribution Models

As the performance and reliability of species distribution models (SDMs) can be affected by sample size^103^, we excluded species with fewer than 13 occurrences since this has been the proposed threshold considered as sufficient for generating reliable SDMs for widespread species^104^. Therefore, only *Vulpes corsac, Lutra lutra and Martes foina* were excluded, as recently developed methods specifically designed for rare species are more appropriate for modeling them^105^. The species included in the SDMs were *Ursus arctos, Ursus spelaeus, Panthera spelaea, Panthera pardus, Felis silvestris, Lynx Lynx, Lynx pardinus*, *Crocuta crocuta, Canis lupus, Cuon alpinus*, *Vulpes vulpes, Vulpes lagopus, Gulo gulo, Meles meles, Martes martes*’ *Mustela nivalis*, *Mustela erminea*, *Mustela putorius*, *Homo neanderthalensis* and *Homo sapiens*.

Moreover, multiple observations from the same site or area and time slice may produce sample biases linked to differential sampling efforts and increase the likelihood of spatial autocorrelation ^106^. Therefore, we kept only one occurrence point per grid cell and time slice for each species. Accordingly, we thinned the occurrence data to eliminate clusters of sites, thereby preventing oversampling of environmental conditions from certain areas and reducing potential spatial autocorrelation. Thus, the minimum distance between any pair of occurrence points was 0.5° (ca. 50 km^2^).

For each species, we generated a set of background points from a convex polygon enclosing all the sites where the species was present. Then, we created a buffer around the polygon with a radius equal to 10% of the maximum distance between actual species occurrences ^40^. This area is referred to as the calibration area. The calibration area was used to ensure that pseudoabsences were geographically distributed based on the distribution and density of the actual occurrence data, thereby preventing their placement in regions lacking species records ^52^.

We extracted, for each species, all the available climate and environmental data at each occurrence and background point. We trained SDMs with four different algorithms: Generalized Linear Model (GLM), Generalized Additive Model (GAM), Maximum Entropy Model (MAXENT), and Bayesian Additive Regression Trees (BART). Following previous studies, for GLM, GAM, and MAXENT, a set of background points was generated by sampling, for each occurrence, 50 random locations matched by time within the calibration area^107^. In contrast, for BART, we used an equal number of pseudoabsences and presences, ran the BART model, and repeated this process 10 times. From these ten BART models, we created an ensemble mean projection^108^.

Whereas GLM assumes a linear relationship between the dependent variable and the covariates, GAM can reveal non-linear effects of the independent variables on the dependent one ^109^. MAXENT is a presence-only modeling method widely employed for extant and extinct species due to its robustness to small sample sizes and its ability to model complex non-linear relationships^110,111^. Lastly, BART is a recently developed machine learning methodology that does not assume any specific functional form for the relationship between predictors and the response^112^. Instead, it models the relationship nonparametrically by using a sum of regression trees, allowing for complex interactions (for details, see Supplementary Note 2). For each algorithm, we evaluated the predictive performance with the Area Under the Curve (AUC) and the Boyce Index (BI). To avoid using poorly calibrated models, only projections obtained from SDMs with AUC values larger than 0.7 were used ^113^.

SDMs provide habitat suitability, favorability or probability of presence. Habitat suitability and probability of presence can be misleading for comparison between species because they are affected by prevalence ^114^. In contrast, habitat favorability values are not affected by prevalence, which allows direct comparison between species and models ^114,115^. Favorability values indicate how much a specific cell/patch fits within the range of habitats that are suitable for the species. In this study, we used habitat favorability to compare the potential spatial distribution between species. Habitat favorability ranges between 0 and 1, with values higher than 0.5 corresponding to areas where the probability of presence is higher than expected by chance. Thus, to compute the geographic range evolution of each species during the MIS3, we defined favorable areas as comprising the total land where habitat favorability was larger than 0.5. To integrate the outcomes obtained from each modeling algorithm into a robust projection, we adopted the weighted ensemble approach proposed by previous authors ^116^. Thus, we weighted the model predictions based on Equation 1:

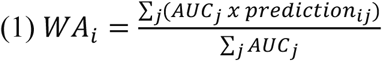

So, for a given species and patch/cell *i*, the weighted average prediction (WA_i_) is calculated as the sum of predictions for site *i* across *j* individual models (GLM, GAM, MAXENT and BART when the AUC values were larger than 0.7) weighted by their respective AUC values and normalized by the sum of all AUC.^116^

Selecting climate or environmental variables is essential for accurate SDMs. However, identifying these variables is particularly challenging for species in the past. To avoid any assumption for the climatic or environmental variables relevant for each species, and to mitigate multicollinearity and overfitting issues, we developed a specific variable selection procedure. First, from the 21 selected variables from HadleyCM3^45^, we removed all correlated variables with correlation coefficients equal to or larger than 0.8. Secondly, we built a function to evaluate, for each species and SDM algorithm, all combinations possible of the remaining variables. For each combination, we used three criteria to select the final covariates: 1) To avoid multicollinearity, all combinations where any of the independent variables had a Variance Inflation Factor (VIF) larger than 5 were excluded; 2) among the combinations where all variables had a VIF <5, we selected the one with the highest AUC value; 3) when more than one combination provided similar AUC values, we selected the combination with the smallest Root Mean Square Error (RMSE). The function performs a stepwise removal of highly correlated variables, prioritizing combinations with the highest AUC and lowest RMSE values. However, the order of variables influences the final selection: general variables such as BIO1 (Mean Annual Temperature) and BIO12 (Mean Annual Precipitation), which appear earlier in the list, are more likely to be retained. This approach is intended to preserve broader climatic signals over more specific or derived variables.

To evaluate the importance of each predictive climate and environmental variable in the SDMs of each species, different specific tests are available according to the model used. Thus, for BART models, the variable importance is commonly measured as the proportion of total branches used for a given variable. In contrast, for linear models the t-statistic can be used to assess variable importance, whereas permutation tests are frequently used for maximum entropy models. In this study, we used these specific methods for calculating variable importance in each model (Extended Data Figure 1). To standardize the variable importance across models and determine the importance of each variable in the final ensemble projection for each species, we adopted the weighted ensemble approach previously described (WA_i_). Therefore, each variable’s importance in the final ensemble model was weighted by its respective area under the curve (AUC) value (Extended Data Figure 1). This method ensured that each climate and environmental variable’s contribution to the final species ensemble projection was proportionally represented.

To further assess the robustness in predicting species distributions, we used block cross-validation. Block cross-validation techniques have demonstrated their efficacy evaluating model transferability, which refers to the capability of extrapolating predictions to novel areas or periods^116^. We used a five-fold spatial block cross-validation resampling method to generate a set of training and test datasets. Therefore, we first randomly split the geographic space into five roughly equal-sized parts, and then fit the models five times. Each time, the occurrence points (i.e., presences) located within one of the blocks was used as test data and the remaining as training data. The block size was determined using the “cv_spatial_autocor” function from the BlockCV R package ^117^, which defines the appropriate block size based on spatial autocorrelation. The subsequent analyses only included the SDMs with good performance and robust model transferability according to the block cross-validation.

From each WA projection, we examined the niche breadth of each species and the degree of niche overlap between pairs of species. To assess niche breadth, we used Levin’s B1 and B2 metrics, reflecting the heterogeneity of environmental conditions where a species is present^118^. To evaluate niche overlap, we computed both Schoener’s D and Warren’s I values, which quantify the similarity extent in the environmental or climatic preferences between two species^119,120^. Additionally, we computed potential spatial overlap by quantifying the number of favorable patches shared by both species at each time slice. We also calculated the geographic range by determining the extent of favorable areas for each species.

To account for age uncertainty, and therefore the uncertainty in the climate values used, we generated a set of 100 SDM replications for each species.^69^ At each replication, we randomly sampled each date from a normal distribution within the 95% confidence interval of the archaeo-paleontological level and used this chronology instead of the calibrated median date to obtain the paleoclimate data. This procedure was repeated one hundred times to assess the extent to which age uncertainty could influence the obtained outcomes. Following previous studies, this sensitivity test assumes dating uncertainty is not randomly nor uniformly distributed within the 95% CI, but follows a Gaussian distribution due to the likelihood structure of calibrated dates, where probability density is highest around the mean estimate and decreases toward the confidence interval limits^121,122^.

### Co-occurrence analyses

Co-occurrence between secondary consumers changes through time.^123^ One of the goals of this study is to evaluate to what extent the transformations in the species distribution ranges and the replacement of Neanderthals by *H. sapiens* affected the co-occurrence patterns among secondary consumers. Therefore, we divided the co-occurrence analyses into four specific periods or phases based on the observed shifts in both the habitat favorability for secondary consumers and the replacement process of Neanderthals by *H. sapiens*. To evaluate whether these phases corresponded with transformations in the richness of secondary consumers in the archaeo-paleontological assemblages, we computed the Shannon index.

To assess the degree to which co-occurring patterns in the fossil record exhibit structured organization, we applied the probabilistic co-occurrence analysis proposed by Veech^124^. We chose this analysis over joint Species Distribution Models (jSDMs) because it provides a straightforward method for measuring species co-occurrence while avoiding the complexities of modeling species interactions. Additionally, although herbivores likely played a crucial role in determining the distribution of carnivores, their representation in the archaeological record is much biased due to human prey selection choices. In contrast, Veech’s formula quantifies co-occurrence directly from presence-absence data without assuming underlying environmental dependencies or species interactions, thereby simplifying parameterization. This analysis determines, for each pair of species, the probability that the observed co-occurrence frequency is significantly greater or smaller than expected. Species found together more often than expected are classified as aggregated, while segregation refers to species that co-occur less frequently than expected by chance. Thus, the observed species presence-absence matrix is compared with randomized matrices to test whether the co-occurrence matrix is structured, with species positively or negatively associated with each other ^124^. This analysis provides co-occurrence probabilities and identifies the specific aggregations and segregations between each pair of species. We conducted this probabilistic co-occurrence analysis for each of the aforementioned phases.

To evaluate whether the number of secondary consumer species in each of the four phases is biased by the available sample size in each phase, we ran a rarefaction analysis. Thus, we compared the observed species richness in each subsample/phase with the expected richness with a simulated sample of 300 archaeo-paleontological assemblages, and a bootstrap method (n=500) was used to obtain the 95% CI of each estimate. According to the rarefaction analysis, the richness of secondary consumers within each phase reached the asymptote. Therefore, increasing the sample size is not expected to increase meaningfully the number of species in any of the phases (Extended Data Fig.8). To further assess the potential impact of different sample sizes across phases on our results, we conducted a sensitivity test. Following previous authors ^39^, we removed the effects of different sample sizes between time intervals or phases by randomly subsampling the species-by-assemblage occurrence matrices to 60 assemblages per phase. Subsequently, we conducted the co-occurrence analyses for each subsample and repeated this process 1000 times to generate 95% CI for the number of aggregations and segregations for each species and phase. Accordingly, this sensitivity test allowed us to estimate the number of aggregations and segregations in each phase independently of the sample size.

### Correlation analyses

To evaluate the impact of niche breadth and overlap on the co-occurrences and the frequency of aggregations and segregations between secondary consumers, we conducted correlation analyses. The purpose of these correlation tests is to assess whether the number of co-occurrences, aggregations, and segregations are influenced by factors such as niche breadth (Levin’s B1 and B2), niche overlap (Schoener’s D and Warren’s I), geographic overlap, or the extent of favorable habitats for the species. To address phylogenetic signal in the data, we computed Blomberg’s K. We used a set of 1,000 phylogenetic trees for extant and extinct mammal species, which capture uncertainties in branching times and topology ^78^, to compute consensus edge lengths and calculate Blomberg’s K. The results show that phylogeny does not significantly influence the SDMs’ outputs variables examined in this study (Supplementary Table 4). Therefore, to account for non-independence between observations (i.e., the same species across different periods), we employed generalized mixed models with a Poisson distribution, as the dependent variables represent count data (e.g., number of occurrences, aggregations, and segregations).

The data and R codes used to perform all of the analyses reported in this manuscript are available from https://osf.io/axyvr/?view_only=874290f77924447a8ed8fdeed72ee4bf

## Data availability

The data collected for this study are available on the Open Science Framework repository (https://osf.io/axyvr/?view_only=874290f77924447a8ed8fdeed72ee4bf).

## Code availability

The code used in this study is available on the Open Science Framework repository (https://osf.io/axyvr/?view_only=874290f77924447a8ed8fdeed72ee4bf).

## Acknowledgements

This research was funded by the European Research Council under the European Union’s Horizon 2020 Research and Innovation Programme (grant agreement number 818299; SUBSILIENCE project; https://www.subsilience.eu). M.V.-C. benefited from a Juan de la Cierva Formación Grant (Ref. FJC2021-047601-I). We thank all of our colleagues from the EvoAdapta group.

## Author Contributions

A.B.M.-A. and M.V.-C designed the study. M.V.-C. recompiled the data. M.V.-C. executed the models and analyzed the data. A.B.M.-A. and M.V.-C contributed to evaluating the outcomes. M.V.-C. led the writing with critical input from A.B.M.-A.

